# Systematic discovery of bacterial symbionts in rumen ciliate protozoa

**DOI:** 10.1101/2025.07.29.667580

**Authors:** Ronnie Solomon, Lina Marcela Botero Rute, Raghawendra Kumar, Ravit Mesika, Ido Toyber, Adi Doron Faigenbaum, Sarah Winkler, Omar Eduardo Tovar Herrera, Shany Gerbi, Deodatta S. Gajbhiye, Tanita Wein, Itzhak Mizrahi, Elie Jami

**Affiliations:** Department of Ruminant Science, Institute of Animal Sciences, Agricultural Research Organization, Volcani institute, Rishon LeZion 7505101, Israel; Department of Life Sciences, Ben-Gurion University of the Negev, Beer-Sheva 84105, Israel; Department of Animal Science, the Hebrew University of Jerusalem, Rehovot 76100, Israel; The Mina and Everard Goodman Faculty of Life Sciences, Bar-Ilan University, Ramat Gan 5290002, Israel; Department of Systems Immunology, Weizmann Institute of Science, Rehovot 7610001, Israel; National Institute of Biotechnology in the Negev, Ben-Gurion University of the Negev, Beer-Sheva, 84105, Israel; The Goldman Sonnenfeldt School of Sustainability and Climate Change, Ben-Gurion University of the Negev, Beer-Sheva 84105, Israel

## Abstract

Microbial interactions are fundamental to global ecological and evolutionary processes, exemplified by endosymbiosis between prokaryotes and single-cell eukaryotes that gave rise to organelles. While such associations remain widespread and ecologically important, the diversity and evolutionary dynamics of intracellular symbioses in many microbial ecosystems remain poorly understood. Here, we uncover a hidden layer of microbial complexity in the rumen ecosystem by identifying multiple endosymbiotic associations between ciliate protozoa and bacteria. Using genome-resolved metagenomics on protozoa enriched rumen fractions, we reveal diverse bacterial genomes exhibiting hallmarks of an obligate intracellular lifestyle. These candidate symbionts span several bacterial phyla and include close relatives of known endosymbionts and parasites of protists as well as previously unclassified or presumed free-living bacterial lineages that likely represent overlooked symbiont specialists. Our findings therefore expand the known distribution of bacterial endosymbiosis, establish the rumen – a key site of global carbon and nitrogen cycling – as a promising model for symbiosis research, and demonstrate the power of our approach to uncover hidden symbiotic associations across complex microbial communities. Overall, our results highlight the ubiquity and evolutionary significance of intracellular symbiosis as a shaping force in microbial ecosystems.

## Introduction

Protists are a diverse group of microbial eukaryotes, inhabiting most environments and playing essential roles in microbial community regulation and ecosystem engineering ^1,2^. Their ability to thrive in diverse environments can be often attributed to their numerous associations with bacteria that span across a wide range of functions such as nutrient cycling and detoxification, protection against pathogens, and many more unique strategies increasing the fitness of host and symbiont ^3–7^. The symbioses between protist and prokaryote can range from facultative, context-dependent relationships, to obligate endosymbiosis, which in some well-established cases have led to the emergence of organelles ^7,8^.

Long-term associations between symbiotic bacteria and eukaryotes often leave genomic signatures within the symbiont genomes. These are most noticeable in obligate intracellular symbionts which typically exhibit distinct traits, setting them apart from even their closest kins^9^. Those traits include reduced genome size and diminished metabolic capabilities due to increased dependency on the host, low GC content, and the emergence and acquisition of genes dedicated to interactions with the host cell ^7,10,11^.

Most insights on the molecular basis for mutualistic interactions between protists and prokaryotes comes from a limited number of well-established interaction models primarily involving free-living protists ^12,13^. In contrast, these interactions remain understudied within animal-associated communities, with research largely focused on termite gut flagellates, specifically the parabasalids and oxymonads ^6,7,14^. Studying protist-bacteria interactions has been constrained by several challenges, which include the difficulty of cultivating and maintaining gut-associated protists, and the time and environment-dependent nature of certain types of interactions ^7,12,15^.

Very few animal gut models exemplify the complex interactions between its resident microbes and the host, such as the one found in the rumen, the fermentative compartment of the pre-gastric chambers of ruminant herbivores. The rumen microbial community, encompassing all domains of life, is responsible for the majority of the carbon and nitrogen uptake of the animal via a large cascade of degradation and cross-feeding of the plant material ingested, making it an ideal model for the study of microbe-microbe interactions ^16–19^. In the rumen, anaerobic ciliate protozoa (ciliophora) represent the main eukaryotic domain and can encompass up to 50% of the microbial biomass ^20–22^. Rumen ciliate protozoa, which belong to two main orders endemic to foregut animals ^23,24^; *Vestibuliferida* and *Entodiniomorphida*, were shown to be involved in various metabolic processes, many of which proposed to be the result of interactions with their prokaryotic neighbors. Studies of metabolic associations between ciliate protozoa and prokaryotes are so far mostly limited to the relationship between protozoa and methane-producing archaea ^25,26^, the methanogens carrying a significant agricultural and environmental impact ^27,28^. However, recent evidence suggests that a much larger breadth of interactions can be found between protozoa and bacteria ^2,29–31^. Studies have noted the physical presence of certain prokaryotes with protozoa, either as ectosymbionts or endosymbionts ^32^, with several studies indicating some bacterial lineages being exclusive or highly preferentially associated with protozoa when compared to the free-living community ^29,33,34^.

In this study, we uncovered novel associations between ciliate protozoa and prokaryotes in the rumen. By enriching the DNA of prokaryotes physically associated with the ciliate community, we identified several genomes from diverse bacterial lineages that exhibit hallmarks of obligate symbiosis. By identifying genes and pathways likely involved in host– symbiont interactions, we provide insights into the molecular mechanisms that may underlie these diverse protozoa-symbiont associations. We further demonstrate the intracellular localization of a candidate symbiont across multiple rumen ciliate species, suggesting a broad host range. Using phylogenomics, our findings reveal that rumen ciliates are host to a variety of symbionts phylogenetically affiliated to ‘symbiont-specialist’ lineages previously characterized in other diverse environments and hosts. These include symbionts from insect hosts, flagellate protozoa in the termite gut, and free-living protists ^35–37^. Those lineages are further shown to be associated with yet uncharacterized genomes spanning across many gut environments, including humans, thus reframing previously assembled genomes from certain lineages in the context of symbiotic, and possibly intracellular lifestyle. Our study thus presents a conceptual and methodical framework for symbiont discovery through which we uncovered a yet uncharted diversity of protozoa bacteria symbiotic interaction in the rumen, some of which likely obligatory and ancient in nature, and provides clues about their diversity, adaptation, function and evolutionary dynamics within this complex ecosystem, positioning the rumen as an emerging model system within gut ecosystems for the study of symbiosis.

## Results

### Identification of multiple putative rumen ciliate protozoa endosymbionts

Prokaryotic DNA associated with protists encompasses only a low proportion of the total DNA, estimated to be between 3–8% (Fig. S1a) ^38,39^. Thus, to enable the characterization of the bacterial communities associated with rumen protozoa, we first established a robust pipeline for enriching and extracting prokaryotic DNA from protozoa-associated samples (Fig. 1a). For that, we leveraged a selective lysis method optimized for enriching bacterial DNA in mixed eukaryotic/prokaryotic samples ^40,41^, which we adapted for use with rumen protozoa. The approach proved highly effective, substantially enriching the prokaryotic DNA fraction in our samples (Fig. S1a).

**Figure 1.**
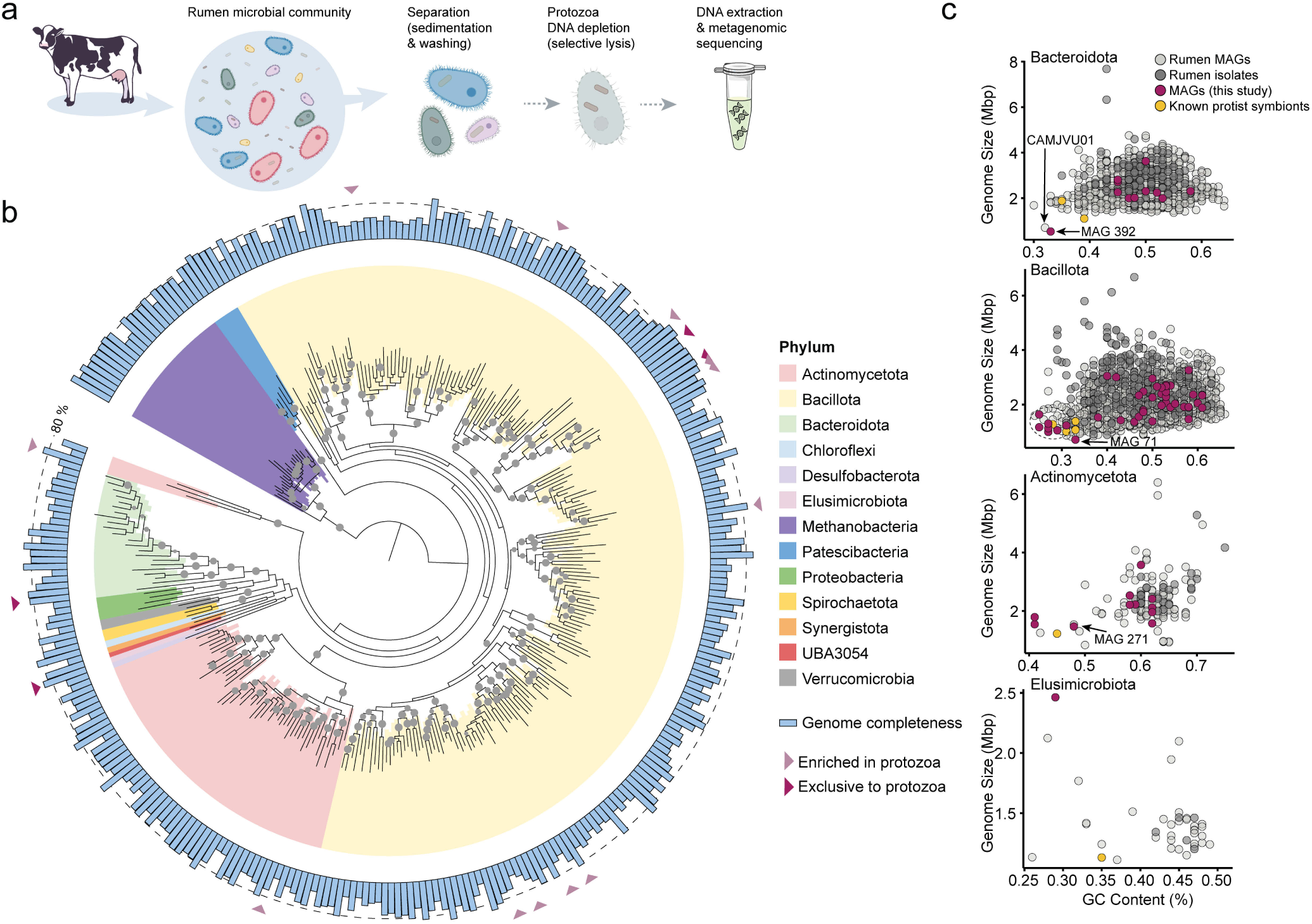
Identification of protozoa-associated genomes and genomic signatures of intracellular lifestyle. **a.** Schematic of the experimental pipeline used to recover metagenome-assembled genomes (MAGs) enriched in the protozoa-associated fraction. **b.** Maximum-likelihood phylogenomic tree reconstructed from 100 single-copy marker genes (SCMG). Branch support values above 75% (1000 ultrafast bootstrap replicates) are indicated as grey nodes. MAGs significantly enriched or exclusively found in the protozoa fraction are indicated in light and dark purple arrows, respectively. **c.** Scatterplot of genome size versus GC content for the high-quality MAGs assembled, alongside curated rumen genomes from large scale metagenomic surveys (light grey) ^43,44^, isolates from the Hungate 1000 project (dark grey) ^50^ and known symbionts from the respective phyla or families if known (yellow). Genomes exhibiting pronounced genome reduction and low GC content, consistent with traits of intracellular symbionts, and further analyzed are indicated.

Subsequently, we set out to examine the prokaryotic genomes associated with rumen protozoa. To this end, we sampled rumen fluids from five milking cows and obtained the protozoa community by separating it from the fractions containing the free-living prokaryotic community as previously described ^24,42^. Briefly, the protozoa fraction underwent several serial washing steps to minimize free-living and loosely associated prokaryotic cells. Then, the protozoa cells were selectively lysed, enriching for intact prokaryotic cells, and the prokaryotic DNA was subsequently extracted (see methods). Shotgun metagenomic sequencing and *de-novo* assembly of the prokaryotic genomes was performed for each sample as well as the respective free-living prokaryotic communities in order to detect genomes uniquely associated with protozoa. Overall, 318 metagenome-assembled genomes (MAGs) were recovered, of which 80 were of high quality (>80% completeness <5% contamination), and 127 of medium quality (>50% completeness <5% contamination) (Fig. 1b, Fig. S1b). When comparing the newly assembled MAGs to the phylogeny of ruminal MAGs obtained from previous large-scale studies ^43,44^, we found that ∼65% (210) of the MAGs assembled here likely represent novel species, based on the Average Nucleotide Identity (ANI) ≥ 95% ^45,46^ (Fig. S1b). When we compared the abundance of the MAGs assembled in the free-living vs. protozoa-associated rumen fractions using read alignment, we found that several genomes were either significantly enriched or detected exclusively in the protozoa-associated community samples (Fig. 1b, Fig. S2a). These genomes, annotated using the Genome Taxonomy Database Toolkit (GTDB-tk; ^47^), include members from the phyla Actinomycetota represented by the family Bifidobacteriaceae, Bacillota including the families Ruminococcaceae, Acutalibacteraceae, and Lachnospiraceae, Bacteroidota represented by the family of Amoebophilaceae as well as Elusimicrobiota with the family Endomicrobioaceae. Notably, these families were previously found to be significantly associated with the rumen protozoa community ^34,42^. Genome analysis revealed that several of the protozoa associated MAGs across the different families carry traits indicative of intracellular lifestyle. Bacterial obligate intracellular symbionts often exhibit unique genomic and functional characteristics stemming from long-term evolutionary changes leading to metabolic dependency between them and their host. Common, and most noticeable among those include streamlined genomes, characterized by a reduction in genome size containing fewer genes as well as a lower GC content ^10,11,48,49^.

To assess these features we applied comparative genomic analysis using solely the high quality assembled genomes (80), with comparison to other and high-quality MAGs assembled from previous sequencing efforts of the rumen ^43,44^, rumen isolates ^50^ as well as representative known intracellular symbionts/parasites of protists belonging to the same phylum or family, if such are known (Fig. 1c).

Our analysis uncovered that several of the genomes in protozoa-enriched samples showed a pronounced deviation from their respective phylum characteristics, having markedly smaller genome size and lower GC content. (Fig. 1c). Within the Bacteroidota phylum, one genome (MAG-392), encompasses the smallest genome assembled in this study (89.5% completeness), with 0.53 Mbp genome size and a GC content of 33% (average genome size of rumen Bacteroidota: 2.55 +/- 0.6 Mbp, GC: 50% +/- 4.6), and was exclusively detected in protozoa associated samples (Fig. S1c). Interestingly, only one previously sequenced Bacteroidota genome (CAMJVU01) from rumen metagenomes out of 2002 curated available genomes from this phylum ^44^, exhibited such similar extreme characteristics, with a genome size of 0.6 Mbp and GC content of 32%. Notably, both genomes belong to the Amoebophilaceae family based on GTDB-tk taxonomic classification, a family from which various intracellular symbionts have been characterized in other environments, including free-living protists ^37,51^ (Fig. 1b,c).

Within the Bacillota phylum, eight genomes were assembled, with genome sizes and GC content which significantly deviated from the phylum average (average genome size of rumen Bacillota; 2.2 +/- 0.59 Mbp, GC: 48% +/- 7.1), with genome sizes between 0.7 - 1.6 Mbps and GC content 25 - 33%. Of these small genomes, four were observed to be enriched or exclusively found in protozoa associated community samples. Interestingly, all but one genome, including those assembled that are not significantly associated with protozoa, belonged to the same family, Ruminococcaceae, representing one monophyletic clade (Fig.1b). We therefore opted to further assess this clade as whole despite some of these genomes not significantly being associated with protozoa. In addition, the smallest genome from Bacillota (MAG-71; genome size = 0.7, GC = 33%) belonged to the Acutalibacteraceae family based on GTDB-tk classification, this family being the only one to our knowledge for which intracellular symbiotic/parasitic lifestyle have been demonstrated within the Bacillota phylum ^35,36^.

One genome from the Actinomycetales phylum was enriched in some the protozoa-associated communities and in some protozoa associated samples was the most abundant genome found, with a genome size of 1.46 Mbp and GC content 48%, thus showing a pronounced deviation from the phylum’s genome size (2.46 +/- 0.76 Mbp) and characteristic high GC content (61% +/- 5.1). Although this genome represented one of the largest genomes among our candidates, its enrichment in the protozoa associated community, along with the fact that it was classified as Bifidobacteriaceae, the same family as *Candidatus Ancillula trichonymphae*, a known protist symbiont from Actinomycetota, led us to include this genome in subsequent analyses. One genome from the Elusimicrobiota phylum was also exclusively found in the protozoa associated community. This phylum encompasses the TG-1 clade, a well characterized lineage of intracellular symbiont of termite flagellates ^52^. However, we opted to preclude this for subsequent analyses as it didn’t exhibit reduced genome size (Fig. 1b).

The distinct genomic features identified in these MAGs along with their preferential association with ciliates are a strong indicator that these bacteria are yet uncharted rumen ciliate protozoa symbionts. This provided a strong foundation for further functional and evolutionary analyses aimed at elucidating the breadth, specialization, and potential origins of protozoa–bacteria symbioses in the rumen.

### Protozoa-associated genomes exhibit functional characteristics of obligate intracellular symbionts

In addition to reduced genome size and low GC content, intracellular symbionts typically exhibit reduced metabolic functions, while increasingly relying on the host for essential processes. They also may acquire or retain functions that are essential for interacting with their host cell ^7,48^. In order to test if this is the case for the small genomes associated with protozoa, we analyzed the functional capabilities of the candidate MAGs of the families belonging to the phylum Bacteroidota (2 MAGs; Amoebophilaceae), Bacillota (8 MAGs; Ruminococcaceae, Acutalibacteraceae), and Actinomycetota (1 MAG; Bifidobacteriaceae) (Fig. 1c, Table S1). We first annotated the gene content, identifying key metabolic pathways and evaluating their completeness. These MAGs were then compared to phylogenetically representative rumen MAGs and isolates, as well as to reference genomes of well-characterized intracellular symbionts and parasites from the same phylum or family if known (Fig. 2, Table S2). We also derived the mean gene-count and pathway-abundance profiles for each family using publicly available rumen datasets ^43,44,50^, to quantify the degree of functional adaptation to an intracellular lifestyle (Fig. 2).

**Figure 2.**
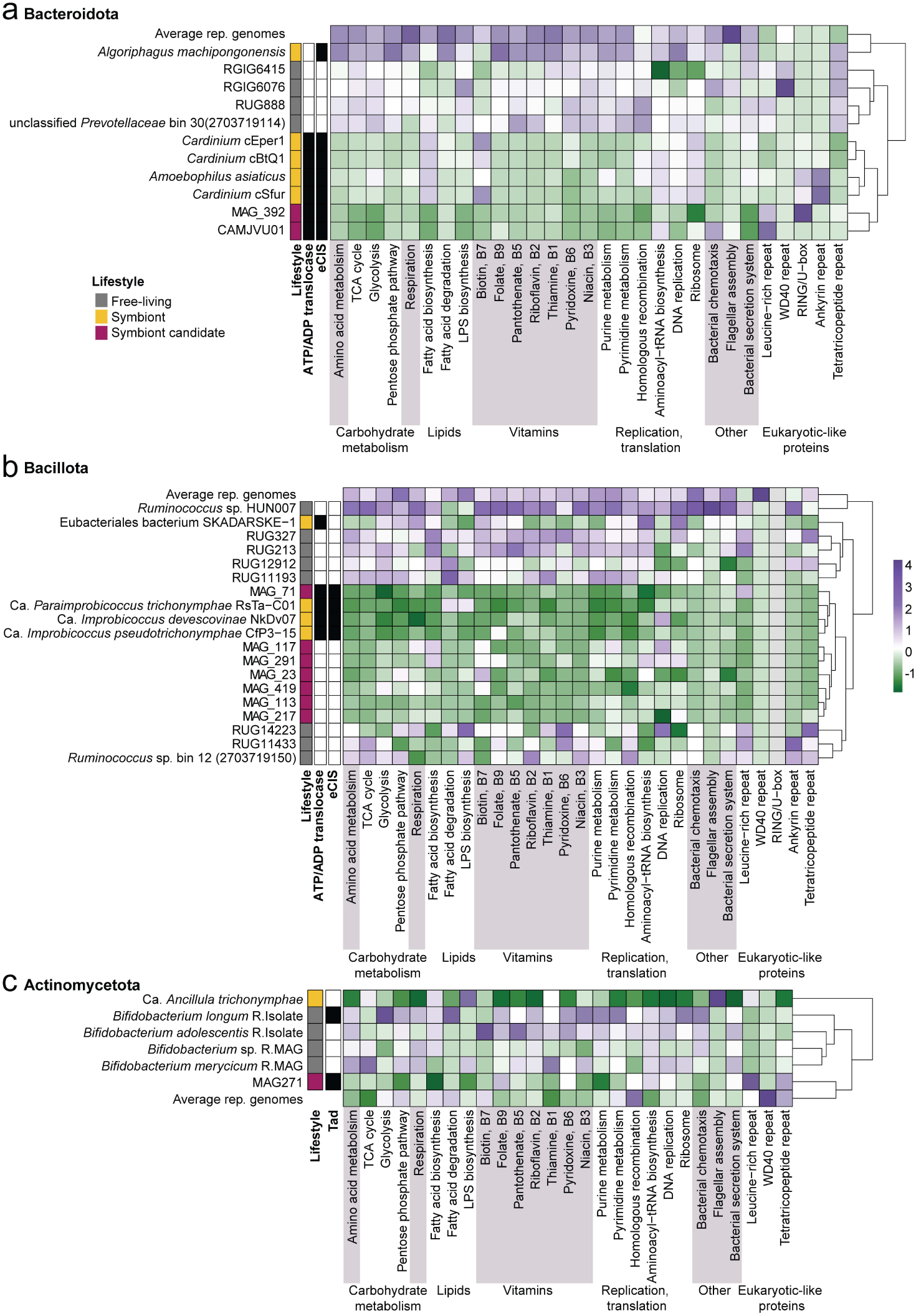
Functional characterization of candidate symbiont genomes. Heatmaps showing the z-score normalized number of proteins in key metabolic processes, pathways and functions across candidate symbionts from the families **a.** Bacteroidota; Amoebophilaceae, **b.** Bacillota; Ruminococcaceae and Acutalibacteraceae and **c.** Actinomycetota; Bifidobacteriaceae, compared to reference genomes from selected MAGs and isolates (lifestyle; grey squares), as well as known intracellular bacterial species (lifestyle; yellow squares) from their respective families. The presence of notable genes such as the ATP/ADP translocase gene, and secretion systems (e.g., eCIS, TAD) in the candidate genomes are indicated in black squares on the left side of the plot.

Our analysis revealed fewer protein-coding sequences (CDS) compared to close relatives of the free-living ruminal genomes in most of the candidate genomes ranging between 500 - 1200 (Table S1). This reduction encompasses central metabolic genes and pathways essential for energy metabolism, carbohydrate metabolism, amino acid biosynthesis, cofactor and vitamin synthesis, replication, and translation, as previously characterized in obligate intracellular symbionts ^11,15,49^ (Fig. 2; Fig. S2). Notably, in the Bacteroidota and Bacillota candidate genomes, the glycolysis pathway is fragmented with key enzymes absent (Fig. 2, Fig. S2, Table S2), impeding the production of ATP. While in the Bacillota candidate genomes, the presence of the phosphate acetyltransferase-acetate kinase pathway (*pta - ackA*) which can generate ATP and may be used to compensate for the absence of the main ATP generating pathway (Fig. S2), no alternative ATP generating pathway could be found in the Amoebophilaceae genomes MAG-392 and CAMJVU01. Instead, to potentially compensate for their lack of energy producing capabilities, these candidate genomes, as well as MAG-71 (Bacillota) encode for the ATP/ADP translocase gene (Fig. 2a,b). This gene which facilitates the import of host-derived ATP across the bacterial membrane is considered a hallmark gene for obligate intracellular bacteria and has been identified in numerous symbionts and parasites across multiple bacterial lineages ^4,15,35,53^. Hence, the presence of the ATP/ADP translocase gene in the putative symbiotic genomes, alongside the reduction of energy-generating pathways, suggests that these bacteria may depend on their ciliate host for energy supply.

Secretion systems, such as type IV or type VI represent well-known components that enable mutualistic, parasitic, or pathogenic interactions with eukaryotic cells ^54–56^. These systems allow bacteria to manipulate host metabolism, evade immune responses, and adapt the host environment supporting bacterial survival ^54,55^. Direct sequence homology search did not uncover any complete type IV and type VI secretion systems in our candidate genomes, however we observed several genes annotated as phage tail proteins. Upon further inspection and combining homology search using a dedicated database^57^, as well as structural similarity, we identified the complete set of genes belonging to the extracellular contractile injection system (eCIS) in the candidate Amoebophilaceae genomes MAG-392 as well as the Acutalibacteraceae MAG-71 (Fig. 2a,b, Table S3). This system, resembling bacteriophage contractile tails, shares multiple genes with type VI secretion systems, and was previously identified in multiple intracellular symbionts ^4,15,58^. This system is suggested to facilitate interactions with host membranes and has been recently implicated in phagosomal escape in the symbiont *Candidatus Amoebophilus asiaticus,* also belonging to the Amoebophilaceae family ^59^. Although carrying genes unique to the known eCIS, the *Candidatus Amoebophilus asiaticus* system was shown to not be extracellularly secreted and thus was defined as T6SS^IV^, encompassing a fourth and distantly related to the canonical type VI secretion system known ^59^.

Similarly to the Bifidobacteriaceae symbiont in termite flagellate, *Candidatus Ancillula trichonymphae* ^60^, the Bifidobacteriaceae rumen candidate symbiont MAG-271, did not exhibit the extensive reduction in core metabolic functions like the other candidates despite having a reduced genome (Fig. 2c, Fig. S2). While no type IV or VI secretion systems were found it did encode for the complete tight adherence (TAD) secretion system (Table S3), known for its role in adhesion to host cells, biofilm formation, and possibly predation evasion, and has been recently associated with episymbiosis ^61^. This system has also been identified in the deep-branching Rickettsiales family Mitibacteraceae, which are hypothesized to represent a transitional stage toward obligate intracellular lifestyles ^56^. This may explain the genome’s high abundance in both the free-living community and in the protozoa associated fraction in which in some samples it encompassed the most abundant genome.

Taking together the reduced genome sizes, low GC content, limited metabolic capabilities, including in some cases the lack of complete ATP-generating pathways, coupled with ATP/ADP translocase genes, and the identification of specialized secretion systems, our results highlight adaptations of multiple of the candidate genomes to obligate intracellular lifestyles.

### Characterizing putative interactions with the host based on eCIS genomic region

The parallel discovery of the eCIS genes in two distant symbiont lineages belonging to different phyla suggests that this secretion system may be a common strategy among symbionts that plays a central role in mediating host-symbiont interactions. In order to understand the evolutionary relationships and potential functional convergence of function in symbionts, we performed a phylogenetic analysis on the phage tail sheath protein belonging to the eCIS locus. This analysis revealed that the genes do not cluster according to the phylum-level taxonomy of the microbes, their environmental origin, or lifestyle. Instead, clustering reflects the eCIS subtype classification, consistent with previous findings ^62^. Specifically, the Acutalibacteraceae symbiont clusters within the subtype IIb group, along with several Acutalibacteraceae members recently proposed to be termite flagellate intracellular parasites ^35^, while the Amoebophilaceae forms a monophyletic clade with *Amoebophilus asiaticus* and *Cardinium* species, all belonging to the Ib subtype. These findings suggest an independent ancient acquisition of this system in different putative rumen ciliate symbionts (Fig. 3a).

**Figure 3.**
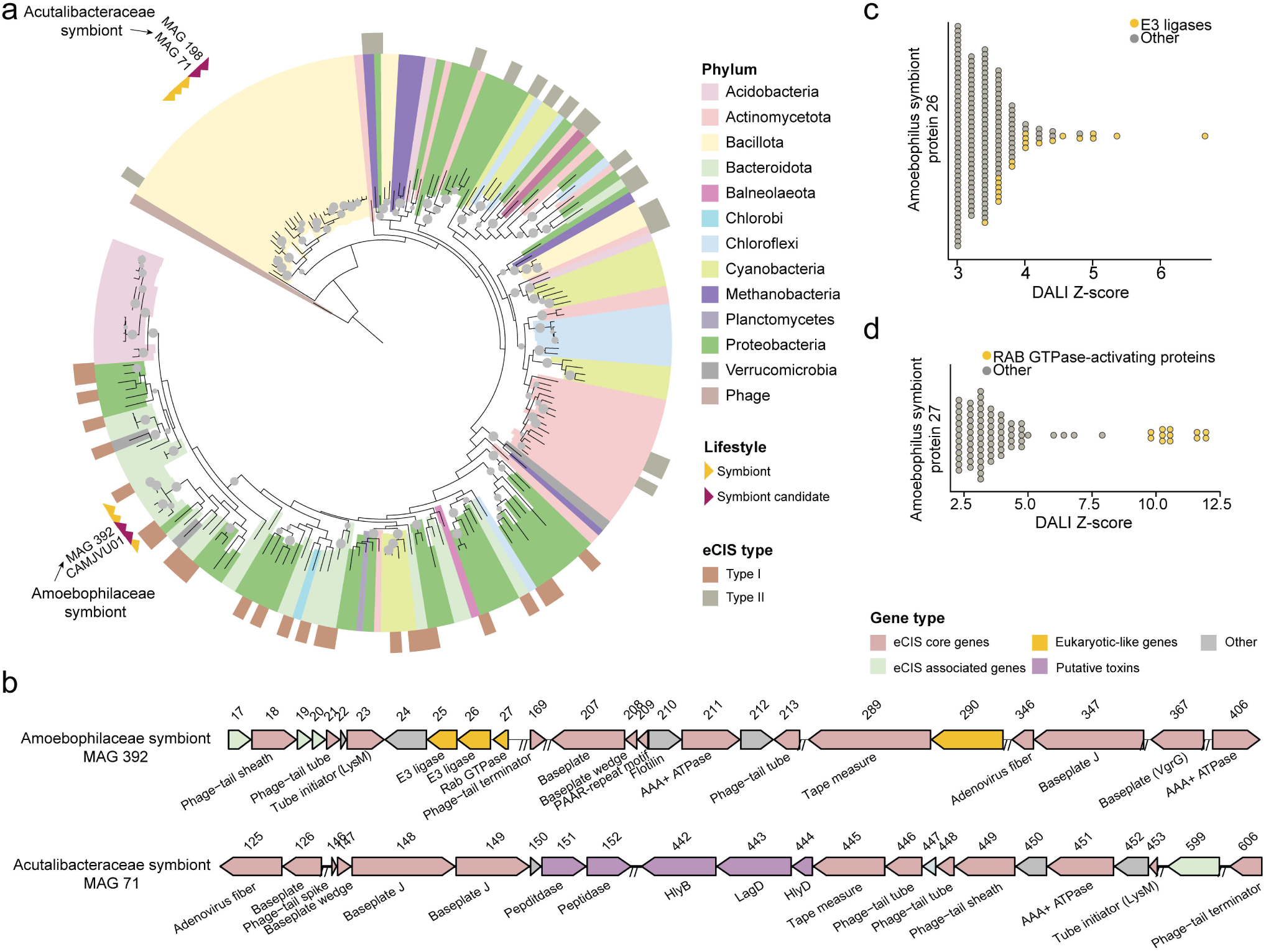
Phylogenetic and organization of the eCIS and associated genes in candidate symbionts MAG-392 and MAG-71. **a.** Phylogeny of the eCIS sheath protein (afp9), showing that the affiliation type of MAG-71 and MAG-392 CIS belong to different subtypes ^57^. Genomes from this study are indicated with a purple arrow and a yellow arrow is used to indicate known symbionts from the same families. **b.** Organization of the eCIS genes in the Amoebophilaceae MAG-392 and Acutalibacteraceae MAG-71 genomes, and adjacent genes encoding for potential accessory functions. **c.** DALI Z-scores of protein structures similar to that of the putative symbiont RING/U-box E3 ubiquitin ligase domain. **d.** DALI Z-scores of protein structures similar to that of the putative symbiont Rab-GAP domain.

Interestingly, when we assessed the prevalence of the eCIS system in the rumen by specifically searching for its genes, we observed that it was found in around 1% of the bacterial genomes (47), and 7 additional genomes from this study. Surprisingly, we discovered that the eCIS system can be found in an additional genome assembled here from the Acutalibacteraceae family (MAG 198, Fig. 3a), and clustering with MAG-71. This genome has not been accounted for due to its relatively low completeness (75%), and no significant enrichment in protozoa associated samples, yet displayed a significant reduction in genome size (0.7 Mbps) as well as low GC content similar to the Acutalibacteraceae MAG-71. Thus, similarly to the Amoebophilaceae clade from the rumen, symbionts from *Acutalibacteraceae* may encompass multiple species associated with ciliate protozoa in the rumen, much like the multiple symbiont strains found in the termite flagellates for the same family ^35^.

While analyzing the genomic neighborhoods of the eCIS systems, we observed that several adjacent genes encoded for genes that could hint at the potential nature or mechanism of interaction between symbiont and host. In the case of the Amoebophilaceae MAG-392, we observed genes encoding for eukaryotic-like proteins (ELPs), proteins which are suggested to mimic host proteins and facilitate host interaction by manipulating cellular processes, evading immune responses to promoting symbiotic persistence ^63–65^. Specifically, genes encoding for RING/U-box, a type of bacterial E3 ubiquitin ligase were found next to eCIS genes, as well as leucine rich repeat proteins (Fig. 3b,c). Such genes and motifs may facilitate host interactions or evasion of host defenses via molecular mimicry ^63^. For example, a E3 ubiquitin ligase in the intracellular pathogen *Shigella flexneri* was recently shown to interact with host proteasome machinery, thereby promoting intracellular persistence ^66^.

The *Amoebophilus* symbiont and its closest relative CAMJVU01 (Fig. 3d), includes other proteins of unknown function containing eukaryotic-like domains, such as a Rab GTPase-activating protein (Fig. 3c). Notably, the symbiont Rab GTPase-activating protein has high structural similarity to the human Rab GTPase-activating protein domain of the RABGAP1L protein (PDB 3HZJ, Fig. 3d) and the TBC domain of human RAB GTPase-activating protein 1 (PDB 4NC6 Fig. 3d). The TBC domain is the common feature among many Rab-GTPase-activating proteins ^67,68^.

In contrast, MAG-71 eCIS system gene neighborhood did not contain ELPs but we identified putative toxins and toxin secretion systems, which could suggest this genome as a potential defensive symbiont. The enrichment of toxins in the genomic neighborhood of the eCIS system has been previously noted with several genes validated for anti-bacterial and antifungal activity ^62^. The gene differences associated with eCIS suggest a different type of association between the two genomes despite the converging strategies for intracellular lifestyle.

Together, these findings highlight how variation in the genetic context of eCIS systems may predict divergent symbiotic strategies, shedding light on the molecular adaptations that shape distinct modes of host-microbe interactions.

### The *Amoebophilaceae* symbiont resides within protozoa cells and clusters with symbiont specialist lineages

We next sought to explore the Amoebophilaceae symbiont (MAG-392) with regards to the physical association with its protozoal host cells as well as its prevalence across different animal habitats. In order to do so, we retrieved the near complete 16S rRNA gene (1200bp) from the assembled genome. We used the 16S rRNA to assess the intracellular location of the Amoebophilaceae *symbiont* within ciliate cells by performing hybridization chain reaction (HCR) fluorescence *in-situ* hybridization (FISH) experiments. For this we hybridized the protozoa cells with a general bacteria probe (red) as well as a probe specific to the symbiont (green, Fig. 4a). We detected the Amoebophilaceae symbiont signal in protozoa species of both the Entodiniomorphida order and Isotrichidae family (Fig. 4a, Fig. S3). Our results confirm that the Amoebophilaceae symbiont is indeed localized within protozoa cells. Our observation further suggests that this symbiont is likely capable of interacting with a broad range of ciliate hosts in the rumen or the existence of multiple strains with different host specificity.

**Figure 4.**
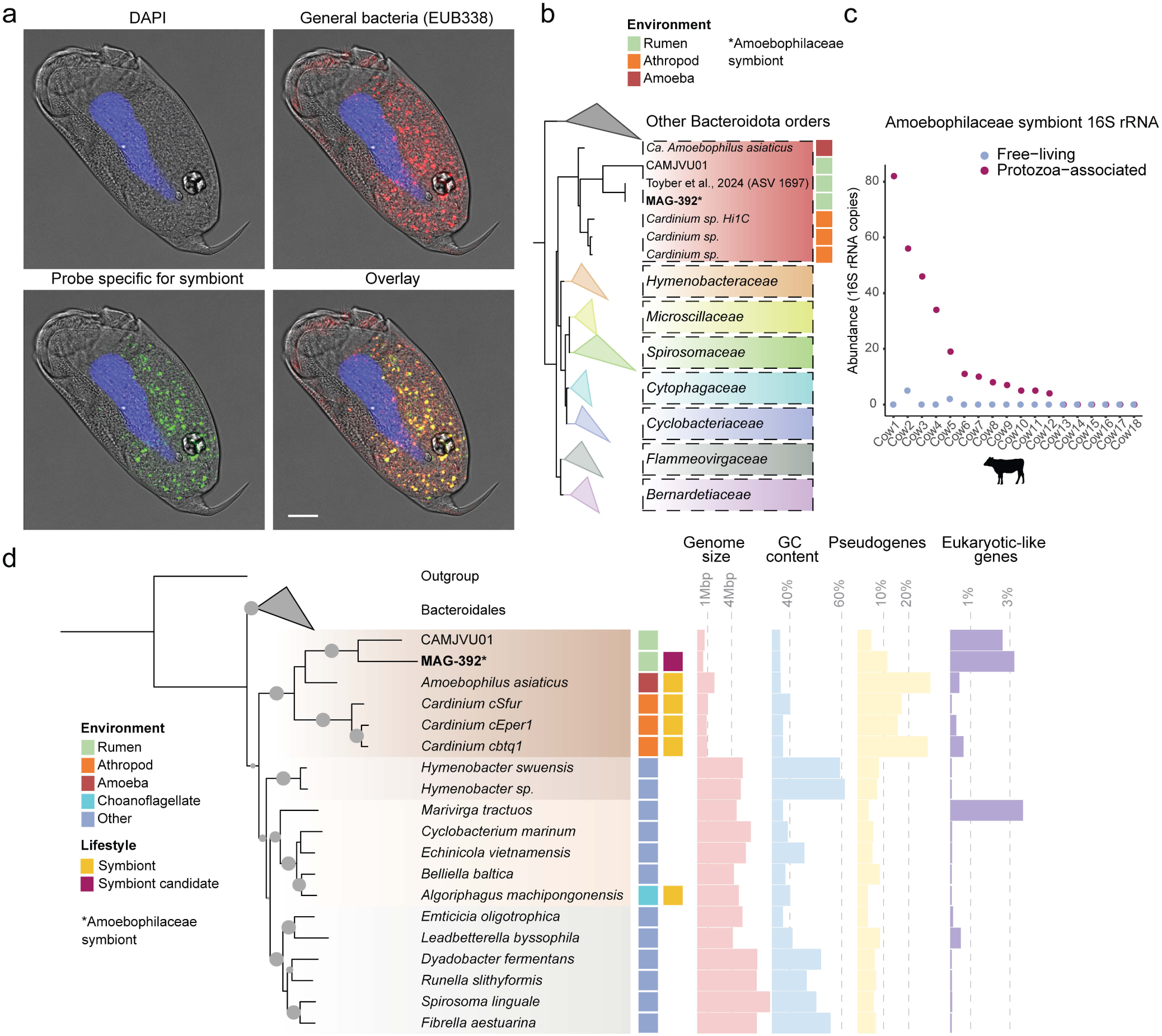
Localization, prevalence, and phylogenetic identity of the Amoebophilaceae symbiont *Ca. Cillophilus ruminis.* **a.** Microscopy images of *Epidinium ecaudatum* cells. HCR-FISH imaging shows intracellular localization of symbiont *Ca. C. ruminis* (labeled in green) within a ciliate protozoa. The total EUB338 bacterial probe is shown in red. DAPI staining shows the macronucleus of the ciliate (scale 10 µm). **b.** Phylogenetic tree of 16S rRNA sequences of Bacteroidota from the SILVA database ^72^ placing MAG-392 within the Amoebophilaceae family alongside other symbionts including the Acantoamoeba symbiont *Ca. Amoebophilus asiaticus* and arthropod symbionts from the *Cardinium* genus. **c.** Distribution and abundance of a 16S ASV identical to MAG-392 16S rRNA across 18 cows ^42^, showing its presence and abundance in the protozoa associated and free-living community. **d.** Genome-scale phylogenomic analysis based on 70 single-copy marker genes confirming the placement of *Ca. C. ruminis* as a novel genus within the Amoebophilaceae.

The retrieval of the 16S rRNA also allowed us to compare its sequence with previous rumen studies in order to assess the prevalence of this genome and its specificity to the protozoa associated community. To do so we first performed a phylogenetic analysis of the of the Amoebophilaceae genome which included Amoebophilaceae sequences from previous rumen surveys and metagenomes available ^42–44^ as well as 16S rRNA sequences from Amoebophilaceae found in other environments (Fig. 4b). The analysis revealed that the Amoebophilaceae symbiont 16S rRNA shared 100% sequence similarity with a single amplicon sequence variant (ASV) from a previous study from our lab, surveying the prokaryotic community composition associated with protozoa (Fig. 4b) ^42^. From this study, we retraced the prevalence of the ASV identical to the one from the Amoebophilaceae symbiont and found that out of 18 cows sampled, the sequence could be detected in 12 cows. In these cows the ASV was exclusively found in association with protozoa while only in two free living samples we detected the ASV in low abundance (below 5 copies, Fig. 4c). These results corroborate the high prevalence of this genome across cows and further demonstrate its exclusive occupancy in the protozoa associated fractions.

Performing a phylogenetic analysis of the symbiont 16S rRNA showed that the Amoebophilaceae symbiont shares 84.7% sequence similarity with *Candidatus Amoebophilus asiaticus,* an intracellular symbiont of free-living Acanthamoeba. It shares high sequence similarity with obligate symbionts from the *Cardinium* genus, a widespread genus, estimated to be in symbiosis with 6-10% of all arthropods, such as *Cardinium hertigii* (Fig. 4B) ^51,69^. The phylogenetic placement of the Amoebophilaceae symbiont 16s rRNA within a family of well-known intracellular symbionts or parasites inhabiting a broad range of eukaryotic organisms strongly supports its classification as an intracellular symbiont ^37^ (Fig 4B).

Consistent with the 16S rRNA-based phylogenetic analysis, phylogenomic analysis using PhyloPhlan3 ^70^, positioned the rumen Amoebophilaceae symbiont alongside *Ca. Amoebophilus asiaticus* with an amino acid identity analysis (AAI) of 49.96, confirming it as its closest known relative aside from the other uncovered rumen Amoebophilaceae genome (CAMJVU01), and suggesting it belongs to at least a new genus (Fig 4C) (Fig S4a) ^71^. These findings show that the uncovered rumen Amoebophilaceae genome belongs to a clade of ‘symbiont specialists’, likely adapted to thrive within various hosts, including free-living protists, insects, and here, as a symbiont of gut-associated ciliate protozoa. Based on their phylogeny and genome sequences, we propose that this genome represents a species within a new genus belonging to the Amoebophilaceae family, which we term *Candidatus Cillophilus ruminis,* and referred to as such hereafter. Moreover, our phylogenetic analysis both at the 16S rRNA level as well as the genome level show a clustering to known characterized symbionts which suggests that such analysis could be used to identify and corroborate other suspected MAGs as potential symbiont specialist lineages.

### Cross-environment emergence of putative endosymbiont clades revealed by comparative phylogenomics

As demonstrated for the *“Ca. C. ruminis”* symbiont, phylogenomic links between newly assembled genomes and symbionts from diverse environments can serve as a powerful approach for discovering novel symbiotic relationships across a wide range of environments and hosts. We thus conducted a phylogenomic analysis incorporating all genomes of the remaining families from the identified phyla containing our candidate symbiont genomes (i.e. Bacillota and Actinomycetota), as well as known symbiont genomes of protists from the respective families if known, to provide a comprehensive evolutionary context.

Our phylogenomic analysis of the Acutalibacteraceae symbiont (MAG-71), encompassed 1455 genomes belonging to the Acutalibacteraceae family. This analysis revealed that MAG-71 belongs to a clade of 41 genomes of predominantly gut-associated origin, which are all characterized by a small genome size and low GC content (Fig. 5a,b). Strikingly, this clade includes all of the currently known intracellular symbionts belonging to Bacillota. These encompass the recently discovered symbionts of flagellate protozoa that reside in the gut of termites, *“Candidatus Improbicoccus pseudotrichonymphae”*, *“Candidatus Improbicoccus devescovinae”*, and *“Candidatus Paraimprobicoccus trichonymphae”* ^35^ (Fig. 4, Fig. S4b). These genomes also represent the closest genomes to MAG-71, aside from the ones originating from the rumen in the same clade, based on amino acid identity. The same clade includes the recently found putative endosymbiont bacteria of the anaerobic free-living amoeba *Pelomyxa schiedtii,* recently shown to form a tripartite interaction with a methanogen and its host ^36^(Fig. 4a). Notably, many of the small genomes in the clade come from non-characterized human gut associated genomes (Fig. 5a). We also observed the presence of eCIS systems in most of the clade further suggesting that this system indeed might be a central and ancient adaptation to intracellular lifestyle in this clade (Fig. 4a, Table S5). This is a similar pattern than the one previously noted in this clade in which the ATP/ADP translocase gene was shown to be conserved across its genomes ^35^. Taken together, our findings suggest that the rumen Acutalibacteraceae symbiont MAG-71 genome, much like *“Ca. C. ruminis”*, belongs to a lineage of symbiont specialists spanning diverse environments, with conserved features of obligate intracellular lifestyle and is likely engaged in a symbiotic relationship with ruminal ciliates.

**Figure 5.**
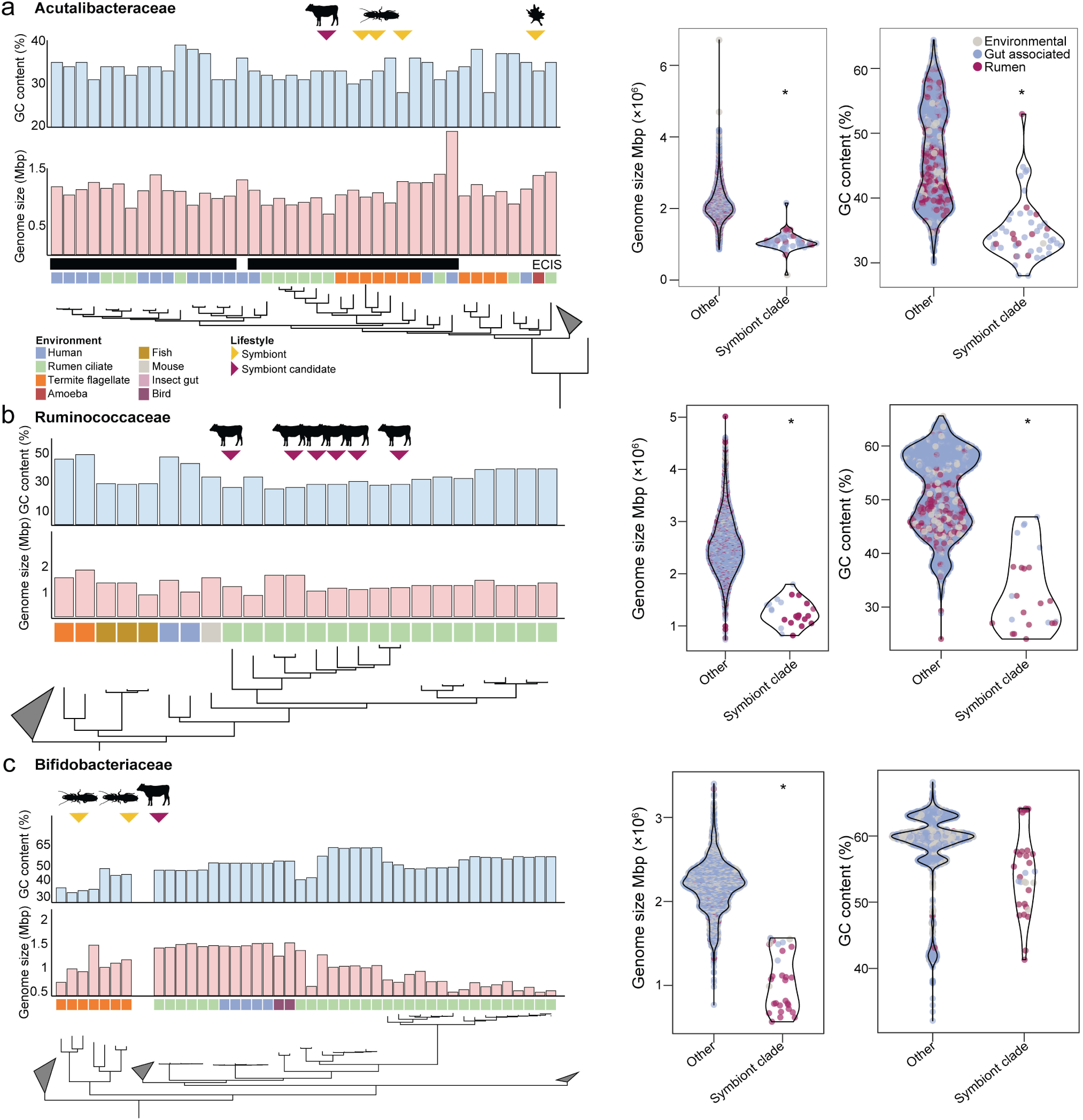
Phylogenomic analysis of symbiont-enriched clades in the rumen microbiome and other environments. Phylogenomic placement of the **a.** Acutalibacteraceae **b.** Ruminococcaceae and **c.** Bifidobacteriaceae candidate symbiont genomes compared to all representative genomes available in the GTDB for each respective family. Purple arrows indicate candidate symbiont MAGs from this study and yellow arrows denote genomes of previously characterized intracellular symbionts in other environments. The colored squares above the tree indicate the origin of the genomes and the pink and blue bars denote the genome size and GC content respectively. The black squares above the **(a)** Acutalibacteraceae tree denote the presence of at least 90% of the core genes of the eCIS (see Table S4). The violin plots next to the trees represent the comparison of the genome size (left plot) and GC content (right plot) of the clade for which the putative symbionts belong against all the unique high-quality genomes found in each family from the GTDB.

We next analyzed the Ruminococcaceae family and its protozoa associated genomes. We performed phylogenomic analysis with these genomes together with all unique representative genomes defined as Ruminococcaceae from the GTDB (1353) ^73^. Our analysis revealed that the Ruminococcaceae symbiont genomes were part of a monophyletic clade of 24 members with other genomes from various sources including rumen, human, termite, fish ^74^ and mice (Fig. 5b). When comparing the average genome size and GC content of the clade to all the Ruminococcaceae genomes available (including those from the current study) we show that their average size is significantly lower than the average Ruminococcaceae genome size and GC content (Fig. 5b, Mann Whitney u test *p* < 0.001). Our analysis therefore indicates that this small genome clade may represent yet uncharacterized symbionts from the Ruminococcaceae spanning across hosts such as rumen, human, fish or termites.

We further analyzed the Actinomycetota genome (MAG-271) associated with rumen protozoa. Phylogenomic analysis of this genome positioned it within the Bifidobacteriaceae family within a clade of genomes exhibiting sizes significantly smaller than the average Bifidobacteriaceae (Fig. 5c). Unlike the Acutalibacteraceae genome, the Bifidobacteraceae MAG-271 assembled here belongs to a distinct clade than the flagellate termite symbionts, *Ca. Ancillula trichonymphae* and the recently proposed *Optitualix* (Fig. 5C). Instead this clade includes several different putative genera, which includes *Scardovia wiggsiae,* a known pathogenic species associated with early childhood caries in humans ^75^. The presence of pathogens in this clade is reminiscent of what has been shown recently in Rickettsiales symbionts of choanoflagellate and their potential connection to human and animal pathogens^76^. The clade is similarly to the other families assessed characterized by mostly small genomes, the average of which significantly smaller than those belonging to the Bifidobacteriaceae family (Mann Whitney u test; *p* < 0.001). In addition, two subclades, defined as RGIG7107 and RGIG1476, with genomes exclusive to the rumen exhibited an even more extreme genome reduction, with some genomes having genome sizes close to 0.5 Mbp.

Thus, anchored by their enriched presence in association with rumen ciliate, our phylogenomic analyses of protozoa-associated symbionts allows to infer that these clades may represent species with symbiotic lifestyle potentially interacting with protists or alternatively in tight association with the host animal cells across different environments.

## Discussion

Our findings revealed a yet uncharted repertoire of microbial symbioses between ciliates and prokaryotes in the rumen. While symbiotic interactions between ciliate protozoa and bacteria are well described in a variety of ecosystems ^7^, no conclusive evidence had previously demonstrated for such associations in the rumen, where ciliates encompass up to 50% of the biomass and belong to lineages exclusive to foregut animals ^23,24^. Here, we show for the first time that rumen ciliates host a diverse and phylogenetically broad array of bacterial symbionts.

Beyond their phylogenetic diversity, the symbionts we identified encode specialized genes and gene clusters that reflect evolutionarily convergent strategies of host adaptation employed across distantly related lineages, which were likely acquired independently. For example, ATP/ADP translocases, which mediate host-derived energy acquisition, are found in multiple lineages and exemplify repeated adaptations to intracellular lifestyles across the tree of life ^4,15,35,53^. Similarly, the presence of the extracellular contractile injection system (eCIS) in two different phyla, belonging to distinct types (I and II), and conserved within their respective lineages, suggests independent, ancient and conserved acquisition and a convergent mechanism facilitating the emergence or maintenance of intracellular interactions. In contrast, we also find clear evidence of lineage-specific host interaction machinery, implying diverse functional roles: putative toxin modules in the Acutalibacteraceae MAG-71 may serve a defensive function, as many eCIS-linked toxins are known to have anti-bacterial or anti-fungal activity ^62^. In comparison, the Amoebophilaceae symbionts designated here as *Candidatus Cillophilus ruminis*, encode multiple eukaryotic-like proteins that suggest direct host manipulation, mirroring what has been observed for its close relative *Candidatus Amoebophilus asiaticus* ^37^. These results suggest that symbiont lineages can share core components of secretion systems yet may diverge dramatically in how they interact with the host and their ecological roles.

The phylogenomic placement of protozoa symbionts within known symbiotic lineages from insects, amoebae, and environmental protists reveals neighboring rumen and other genomes as potential symbionts. What were previously regarded as unclassified or free-living bacteria may, in fact, represent overlooked ‘symbiont specialist’ lineages. The expansion of these “professional symbiont” lineages – from the rumen to free-living protists, termite gut protozoa, and even metazoan hosts – provides a rare opportunity to trace how bacterial lineages repeatedly adapt to intracellular life across disparate environments. For example, the Amoebophilaceae family is remarkable for spanning associations with metazoan cells (*Cardinium* spp.), free-living amoebae (*Candidatus Amoebophilus asiaticus*), and gut-associated protists (*Candidatus Cillophilus ruminis*, CAMVUJ01). Our results therefore underscore the power of genome-resolved phylogenomics not only to resolve evolutionary relationships but also to uncover hidden symbiotic interactions across complex microbial communities, similarly to what has been proposed in the context of endosymbionts phylogenetically associated with intracellular pathogens ^76^. By anchoring these analyses to physical associations with protozoa hosts, as demonstrated here for rumen ciliates, we can expand the known diversity of symbionts beyond classical groups such as the Rickettsiales and Holosporales ^76,77^, and identify entirely novel lineages engaged in host-associated or endosymbiotic lifestyles. Our findings reframe the rumen not merely as a site of metabolic cross-feeding among free-living microbes, but as a dynamic ecosystem shaped in part by previously unrecognized intracellular partnerships. Such interactions likely play critical roles in host biology and microbial community structure, with implications for nutrient flow, energy capture, and possibly the evolution of microbial complexity in the gut, reinforced by previous observations of specific metabolic genes enriched in the ciliate protozoa prokaryotic community ^42^. Understanding how these symbionts influence ciliate ecology and rumen function will be essential for refining predictive models of the microbiome and for advancing sustainable livestock management.

By establishing the rumen as a new and tractable model for studying protist-bacteria symbioses within gut environments, which was previously exemplified primarily by the termite systems, our results enable comparative approaches that span host taxa, symbiont lineages, and ecological contexts. Our study provides a framework for detecting and identifying symbionts across ecosystems. Moreover, this exploration further highlights the power of investigating understudied symbiotic systems to uncover novel principles of host-microbe interaction, shared and divergent ecological adaptation, and the evolutionary forces shaping complex microbial communities. Our findings not only expose a hidden layer of symbiotic diversity in the rumen but also show that these types of interactions are more widespread in gut ecosystems than previously acknowledged and may be relevant to many gut systems. We further suggest that our results call for a systematic reevaluation of microbial associations across animal microbiomes, particularly in environments where protist–bacteria interactions have been historically overlooked, including humans. Recognizing and decoding this complexity will be essential for developing a more complete framework for gut ecosystem function—and for harnessing microbial partnerships in agriculture, ecology, and beyond.

## Materials and Methods

### Animal handling and sampling

The experimental procedures used in this study were approved by the Faculty Animal Policy and Welfare Committee of the Agricultural Research Organization Volcani Research Center approval no. 889/20 IL, by the guidelines of the Israel Council for Animal Care. For the experiment, five cows were fed before the beginning of the experiment under the same high-fiber diet (70% roughage and 30% grains), which is the standard diet for cows during the dry period in our institute.

### Rumen sampling and protozoa community Isolation

Rumen sampling and protozoa community isolation were performed as previously described^34^. Briefly, rumen fluid was collected 2 hours after morning feeding using a custom-made stainless-steel stomach tube connected to an electric vacuum pump (Gast, Inc., MI, USA). Protozoa were counted and characterized in the whole rumen fluid, while the free-living (FL) prokaryotic community was isolated by centrifuging a subset at 500 g for 5 minutes at 4°C.

The absence of protozoa in the FL fraction was confirmed by microscopy. For protozoa-associated prokaryotic community isolation, 1L of rumen fluid was filtered through eight layers of cheesecloth, dispensed into CO₂-filled bottles, and transferred to an anaerobic glove box. The protozoa were sedimented by mixing with anaerobic Coleman salt buffer (1:1) and incubated for 50 minutes at 39°C, followed by glucose addition (1 g/L) for 20 minutes to enhance flocculation. The protozoa were washed four times by centrifugation (500 g for 5 minutes), with buffer replacement, and the final protozoa pellet was suspended in the extraction buffer, frozen at −20°C.

### Enrichment of protozoa-associated prokaryotes

After the isolation of the protozoa community along with their associated prokaryotic cells, enrichment of symbiont DNA was performed by two kits based on selective lysis of eukaryotic DNA on each sample: QIAmp DNA microbiome kit (QIAGEN, Hilden, Germany) and HostZERO microbial DNA kit (ZYMO RESEARCH, Orange, CA, US). Procedures were performed according to the manufacturers. The usage of the two kits allowed for an increase in the number and taxonomic diversity of MAGs assembled. For all the analyses related to the enrichment of MAGs in the protozoa associated samples, the data obtained from the samples that underwent HostZERO depletion were used as we observed that these were more representative of the diversity of the samples based on 16S rRNA amplicon sequencing (data not shown).

### DNA extraction

DNA extraction was performed on the protozoa-associated prokaryotic and free-living communities as previously described ^34^. In brief, cells were lysed by bead disruption using a Biospec Mini-Beadbeater-16 (Biospec, Bartlesville, OK, USA) at 3000 RPM for 3 min with 500ml of phenol followed by phenol/chloroform 1:1. The samples were then precipitated with 0.6 volume of isopropanol and resuspended overnight in 50–100μl TE buffer (10 mM Tris-HCl, one mM EDTA), then stored at 4°C for short-term use or archived at −80 °C.

### Illumina shotgun sequencing

Sequencing was conducted on the eukaryote-depleted protozoa-associated prokaryotic and free-living communities and performed at the Genomics and Microbiome Core Facility, Rush University (Chicago, IL). Illumina libraries were prepared following the Illumina DNA Prep Reference Guide protocol. Briefly, 4 ng of input DNA was fragmented, followed by a post-fragmentation cleanup and 12 cycles of amplification incorporating unique dual indexes to each sample (Nextera DNA UD Indexes). A preliminary pool was generated by mixing equal volumes of each library. The preliminary library was cleaned up with SPB beads and eluted in water. The eluted library was quantified by Qubit and evaluated by TapeStation (D1000 Screen Tape). The resulting concentration was used to load an Illumina MiniSeq Mid-Output reagent cartridge (300 cycles). Sequencing results from the preliminary pool generated a pool representing each sample at equimolar amounts. The new pool was cleaned up with SPB beads and eluted water. The concentration was calculated after the Qubit and Tapestation4150 run. Then, the library was loaded into a NovaSeq6000 2x150 SP lane (56% of the lane).

### MAG assembly, phylogenetic analysis, and genomic and metabolic Characterization

The read adaptors were trimmed by Trim Galore software ^78^, and passed through the quality control process of FastQC. Genome assembly was done according to the MetaWRAP pipeline ^79^. Briefly, contigs were constructed using Megahit version 1.1.3. [N50 > 1050bp]. Binning was performed using three binning software: Metabat2, Maxbin2, and Concoct ^80–82^. Bin consolidation was performed using the bin refinement module in MetaWRAP ^79^. The Quant bin module in MetaWRAP, which utilizes the Salmon tool, calculates the number of reads assigned to MAGs and computes the abundances ^83^. LEfSe [Linear discriminant analysis Effect Size] was used to find MAGs that are significantly enriched in protozoa-associated samples ^84^. Average amino acid identity AAI were calculated using (http://enve-omics.ce.gatech.edu/aai/), between our MAGs of interest and selected reference genomes and MAGs.

The quality of all assembled MAGs was assessed using CheckM2 ^85^, and only MAGs with a maximum of 5% contamination and at least 30% completeness were included in further analysis. Phylogenetic trees, including all MAG trees in the study, were reconstructed using PhyloPhlAn3 based on amino acid sequences (which were created using the Prokka tool) ^70^. The tool utilized the supermatrix_aa configuration file and the PhyloPhlAn3.0 database.

Taxonomic classification was performed using the GTDB-Tk classify workflow following default settings. This pipeline included genomes compared to the GTDB reference database using Mash and refined with skani; those with high ANI to reference genomes were classified directly. Remaining genomes underwent gene prediction with Prodigal and identification of bacterial or archaeal marker genes via HMMER. Marker genes were aligned, concatenated into a ∼5,000 amino acid alignment, and placed into the GTDB reference tree using pplacer. Final taxonomic assignments were based on phylogenetic placement, relative evolutionary divergence, and ANI. All phylogenetic trees were visualized in iTOL ^86^.

For the GC content and genome size analysis, in addition to our assembled MAGs, MAGs from previous metagenomic studies and isolated genomes from the Hungate 1000 catalog of reference genomes from the rumen microbiome were included ^43,44,50^. Known symbionts from the phylum or if existing from the same family as the candidates were also incorporated (e.g., in the Bacteroidota analysis, where Amoebophilaceae members from other environments were included due to their close relation to the uncovered MAG), along with MAGs/genomes from other environments). All genomes were dereplicated using the dRep tool with default settings with ANI set at ≥ 95% ^87^. This approach was also applied to the MAGs and genomes included in the phylogenetic trees.

Prokka was used to identify coding sequences (CDS) ^88^, which were then analyzed using the InterProScan tool ^89^ to classify proteins into families and predict domains such as eukaryotic-like proteins (ELPs). Known ELPs were assessed ^65^, and the percentage of ELPs occurrences was calculated relative to the total number of coding sequences (CDS). Pseudogenes were identified using the tool Pseudofinder with the annotate function, using the nr database as a reference ^90^. The ARAGORN tool was used to search for tRNA ^91^.

The presence and completeness of metabolic modules across the assembled MAGs, known symbionts, and reference genomes were assessed using the MicrobeAnnotator tool ^92^, which integrates various databases, including KEGG Orthology (KO), Enzyme Commission (E.C.), Gene Ontology (GO), Pfam, and InterPro.

### Identification and characterization of the contractile injection system (eCIS) in the genomes assembled

The protein sequences of MAGs were compared against the eCIS database ^57^, a comprehensive resource summarizing identified potential eCIS loci across a wide range of bacterial genomes. Only results with a p-value < 0.00001, high coverage (above 80%), and genetic proximity to other system-related proteins were considered candidates. A cassette was classified as a candidate for a Contractile Injection System only if it contained several distinct core system proteins. We augmented the search for eCIS genes by performing a structure based homology search for hypothetical genes adjacent to eCIS genes to uncover more eCIS genes as well as functions associated with the system using HHpred, AlphaFold 3.0 and Foldseek ^93–95^. For further protein annotations, the predicted structures were further analysed using the DALI webserver ^96^ .

Phylogenetic analysis of the sheath protein, homologous to afp9 was performed, similar to what was previously done in ^59^. Protein alignment was performed using MUSCLE ^97^, and the module for tree construction was optimized with ModelTest-NG, as described earlier. The phylogenetic tree was constructed using raxml-ng with maximum likelihood and bootstrap analysis.

Protein sequences from the genome were locally searched against the Toxinome database ^98^, a comprehensive bacterial toxin database, using the DIAMOND tool to identify putative toxins associated with the system. Successful hits were determined based on the default parameters of the web platform, which included a minimum identity of 40%, a minimum query coverage of 90%, a minimum reference coverage of 60%, and a maximum e-value of 0.001. Toxins were considered potentially related to the system if located near the system within the genome.

### 16S rRNA extraction phylogenetic analysis and MAG-392 ASV quantification

The Barrnap tool was used to extract the 16S rRNA sequences from the MAGs assembled and used to construct a phylogenetic tree for MAG-392 [https://github.com/tseemann/barrnap]. Sequence alignment was performed using MAFFT v7.526 with default settings ^99^. A maximum-likelihood tree was then constructed using IQ-TREE ^100^, which automatically selected the optimal substitution model. Bootstrap support with 1000 replicates was conducted to assess branch support. To construct a phylogenetic tree, representative 16S rRNA sequences from the Cytophagales order were retrieved from the NCBI database as well as identified Cytophagales ASVs from the rumen from a previous study in the lab ^42^.

The ruminal ASV found to be identical to the sequences extracted from the Amoebophilaceae MAG-392 was retrieved and occurrence and abundance across the different samples and fractions (free-living / protozoa associated) was assessed ^42^. The Wilcoxon Signed-Rank test was applied to assess significance between the fractions.

### Fluorescence *in situ* hybridization

To design HCR-FISH probes for the Amoebophilaceae candidate symbiont MAG-392 (*Ca. Cillophilus ruminis*) the full 16S rRNA retrieved from the MAG was used to design 20 probes through Molecular Instruments, Inc. (Los Angeles, CA). The FISH procedure followed their HCR RNA-FISH protocol ^101^. Briefly, protozoa samples were fixed in 4% PFA and dehydrated in ethanol. Pre-hybridization was performed using a hybridization buffer, followed by overnight probe hybridization at 37°C with 2 pmol of each probe. Excess probes were removed through multiple washes with probe wash buffer and SSCT. Amplification was completed by incubating samples in the hairpin solution overnight at room temperature, with subsequent SSCT washes and DAPI staining. Samples were stored at 4°C, protected from light, until microscopy analysis.

## Data Availability

Data that support the findings of this study are available within the article and its Supplementary tables. Both the raw sequencing data as well as the MAGs will be available in the NCBI database upon publication.

## Acknowledgments

We want to thank the ARO farmers and veterinarians for their support throughout all experiments. E.J. was supported by grants from the Israeli Dairy Board Foundation (Grant No. 362-0680) and the Israeli Science Foundation (Grant No. 603/20). I.M. was supported by grants from the Israeli Science Foundation (Grant No. 979/25) and the European Research Council (ERC 866530 and ERC-POC 01082166). T.W. is the Incumbent of Philip Harris and Gerald Ronson career development chair and was supported by a research grant from the Center for New Scientists at the Weizmann Institute of Science.

## Author contribution

The study was designed by R.S., L.M.B.R. and E.J. R.S. performed sampling, metagenomic analyses and phylogenomic analysis. L.M.B.R. performed FISH experiments and phylogenomic analysis. I.T. helped with animal sampling. D.S.G. helped with FISH experiments. A.D.T, S.W., O.T., S.G., and T.W. helped with data analysis. R.S., L.M.B.R. T.W., I.M. and E.J. conceptualized and wrote the manuscript. E.J. supervised the study.

## Notes

### Competing Interest Statement

The authors have declared no competing interest.

