## Supplementary material for "Systematic discovery of bacterial symbionts in rumen ciliate protozoa": Supplmentary figures

### Supplementary Figures

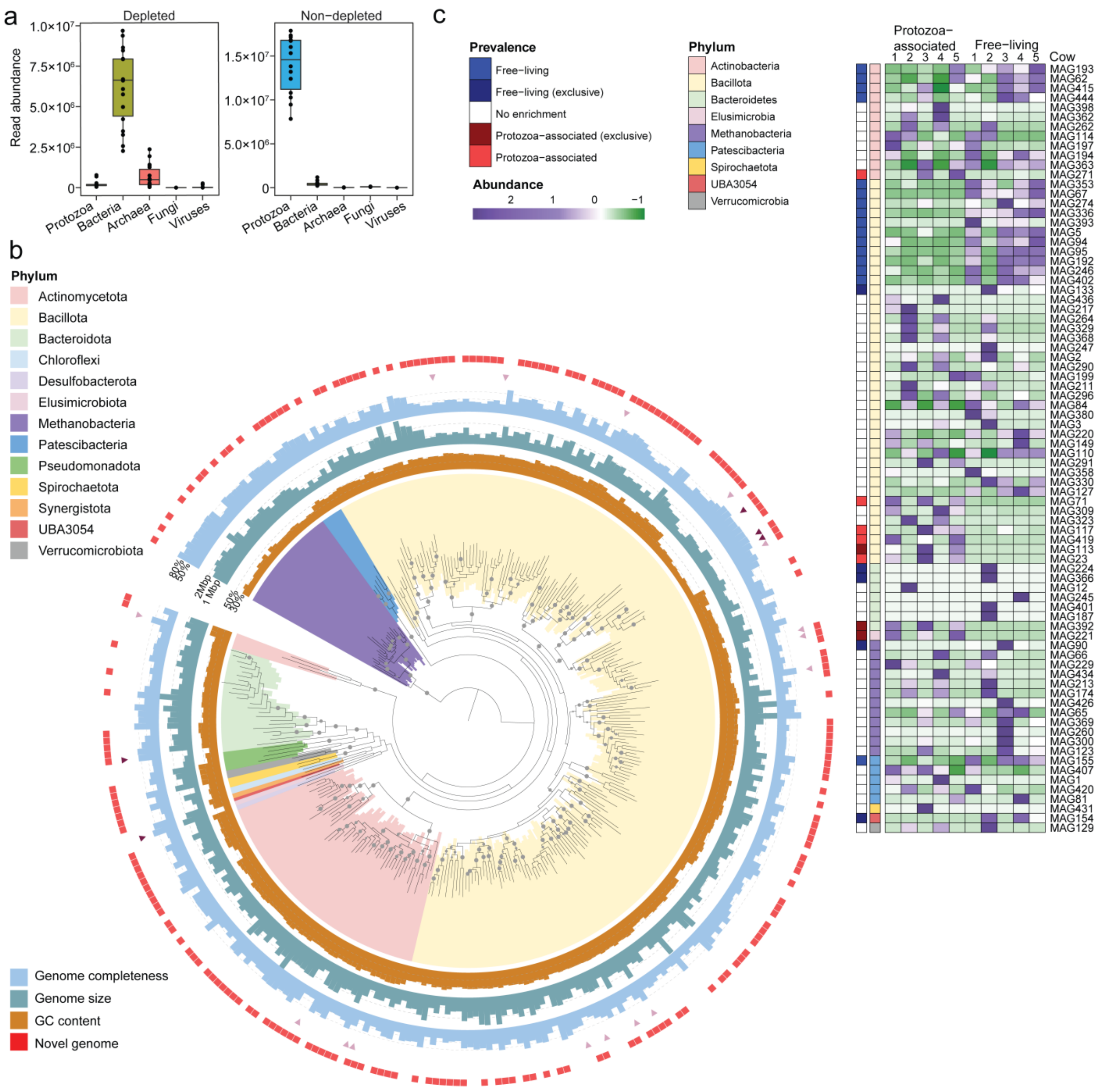


**Figure S1. Enrichment of protozoa-associated symbionts and genome characteristics of assembled MAGs. a.** Comparison of sequencing read origin in samples with and without selective lysis of protozoa. Samples that underwent selective eukaryotic lysis show a marked enrichment of bacterial DNA, overcoming the challenge of low prokaryotic DNA yield in protozoa-associated rumen samples. **b.** Phylogenomic tree of the 308 metagenome-assembled genomes (MAGs) recovered from this study containing all 70 unique PhyloPhlAn marker genes. Inner, middle, and outer rings display GC content (orange), genome size (green), and genome completeness (blue), respectively. Red squares indicate MAGs representing novel species based on Average Nucleotide Identity (ANI < 95%) compared to previously published rumen genomes. MAGs enriched in protozoa-associated samples are indicated by light purple arrows; MAGs exclusively found in the protozoa-associated fraction are denoted by dark purple arrows. **c.** Heatmap of the relative abundance of high-quality MAGs (>80% completeness, <5% contamination) across protozoa-associated and free-living rumen fractions. Rows represent individual MAGs, grouped by phylum (inner colored strip), and columns represent individual samples. The outermost annotation bar indicates MAGs enriched or uniquely found in the protozoa-associated community.

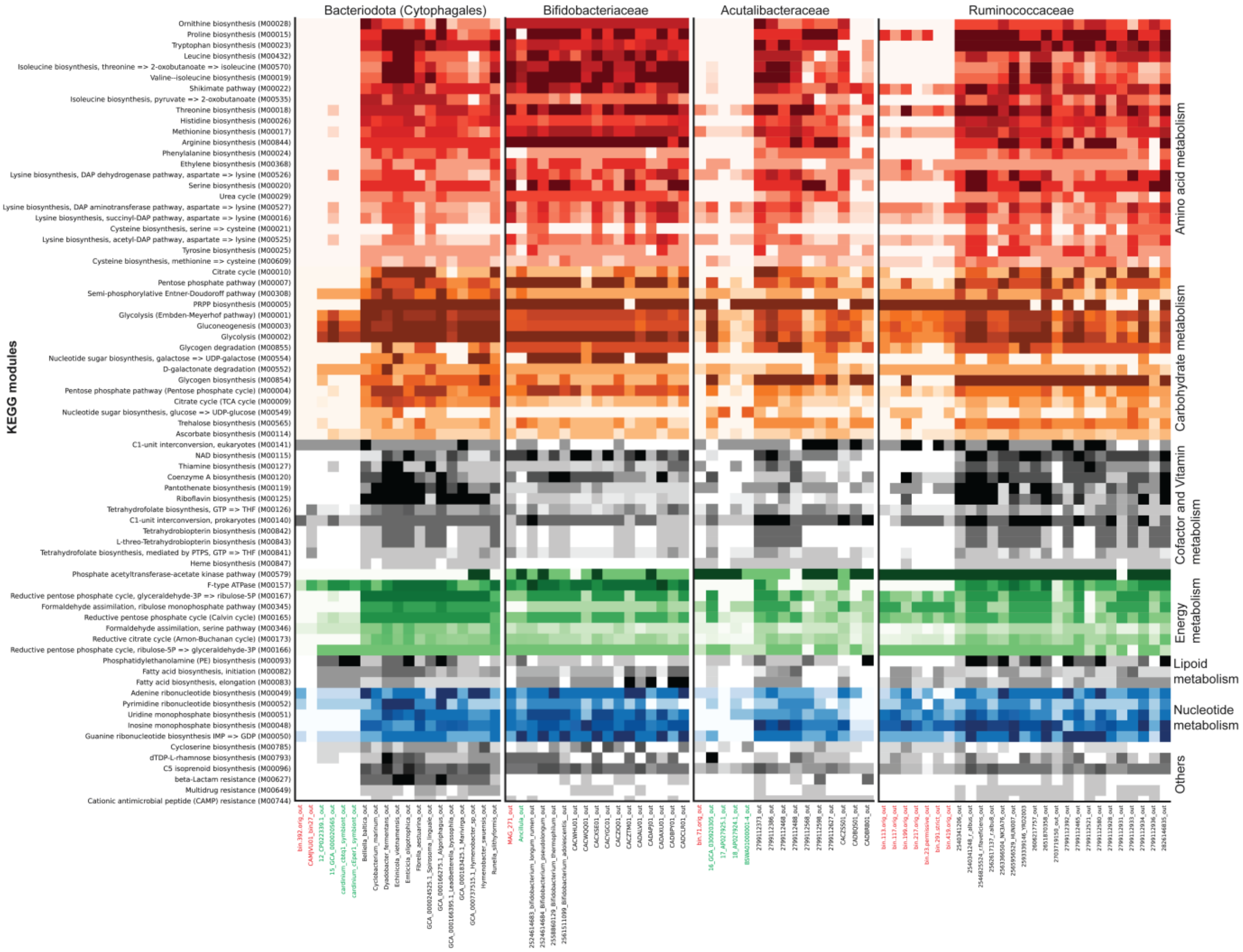


**Figure S2. Detailed metabolic pathway completeness of candidate symbionts and related genomes.** Heatmap showing the percent completeness of KEGG pathways across candidate symbiont genomes and phylogenetically related reference genomes. Pathways are grouped by functional categories, including energy metabolism, carbohydrate metabolism, amino acid biosynthesis, cofactor and vitamin biosynthesis, and core cellular processes. Candidate MAGs identified in this study are highlighted in red, while genomes of known intracellular symbionts from the same taxonomic lineages are shown in green. Darker shading indicates higher pathway completeness. The figure highlights widespread metabolic reduction in the candidate symbionts, with major losses in glycolysis and biosynthetic capabilities, particularly among members of the Bacteroidota (Amoebophilaceae) and Bacillota (Acutalibacteraceae, Ruminococcaceae) lineages. Several MAGs encode the ATP/ADP translocase gene instead of complete ATP-generating pathways, suggesting dependence on host-derived energy. In contrast, the Actinomycetota MAG (Bifidobacteriaceae) shows greater metabolic retention despite genome reduction. Only pathways shared across at least two bacterial lineages are shown here; a comprehensive module list is available in Table S2.

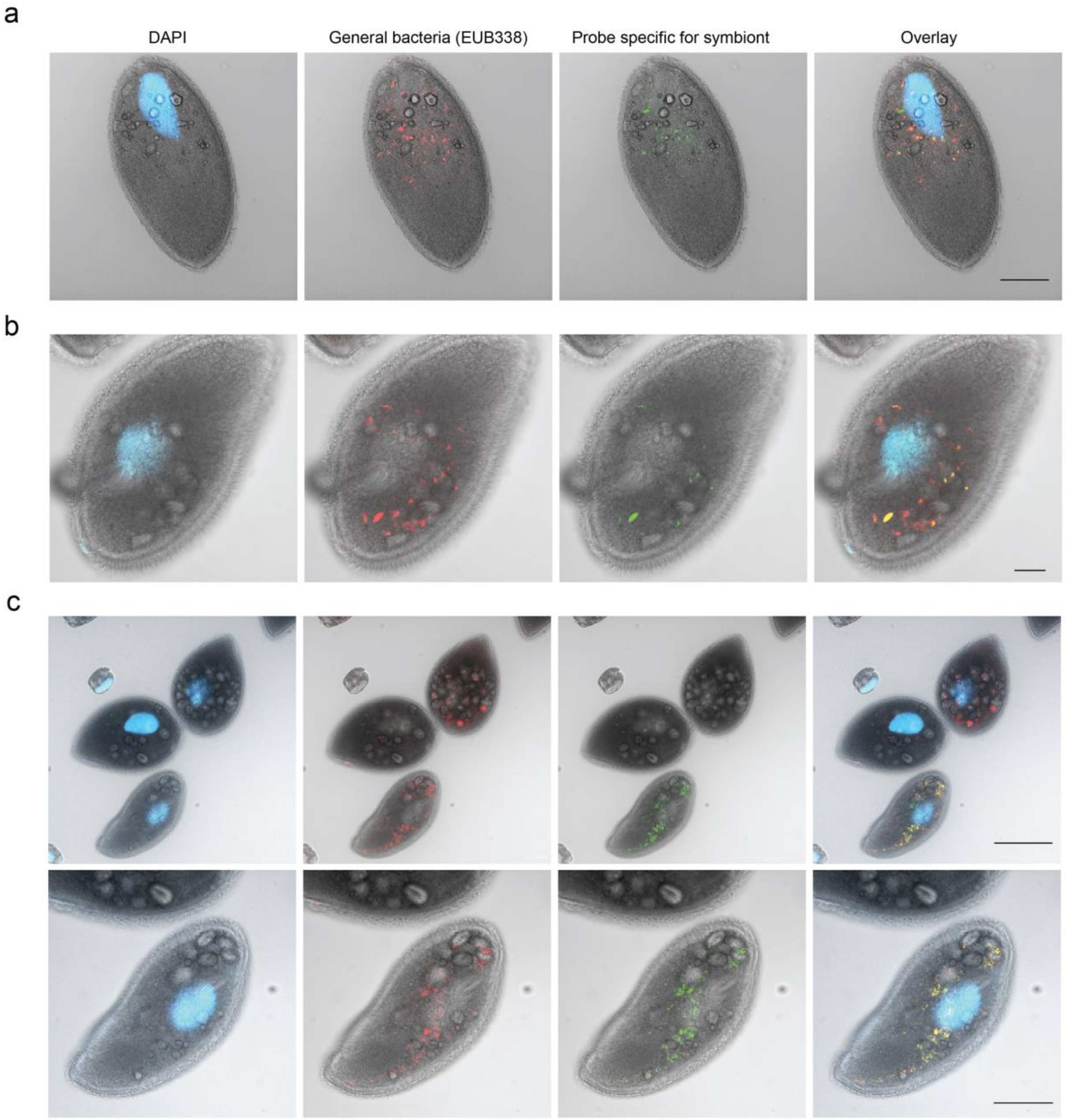


**Figure S3. Localization of MAG-392 within *Isotricha* spp. by HCR-FISH targeting the 16S rRNA.** Hybridization chain reaction–fluorescence in situ hybridization (HCR-FISH) was performed on rumen protozoa using a general bacterial probe (EUB338, red) and a probe specific to MAG-392 (green), both targeting the 16S rRNA. Host nuclei were stained with DAPI (blue), marking the macronucleus of the ciliate protozoa. **a, b.** *Isotricha intestinalis* cells showing intracellular localization of MAG-392. Scale bars: **(a)** 50 µm; **(b)** 20 µm. **c.** FISH imaging of two *Isotricha* species: *I. prostoma* (two upper cells) and *I. intestinalis* (lower cell). While both species harbor bacteria, the MAG-392-specific signal was detected exclusively in *I. intestinalis*. Lower panel shows a magnified view of the *I. intestinalis* cell from the upper panel. Scale bars: upper panel, 100 µm; lower panel, 50 µm.

a

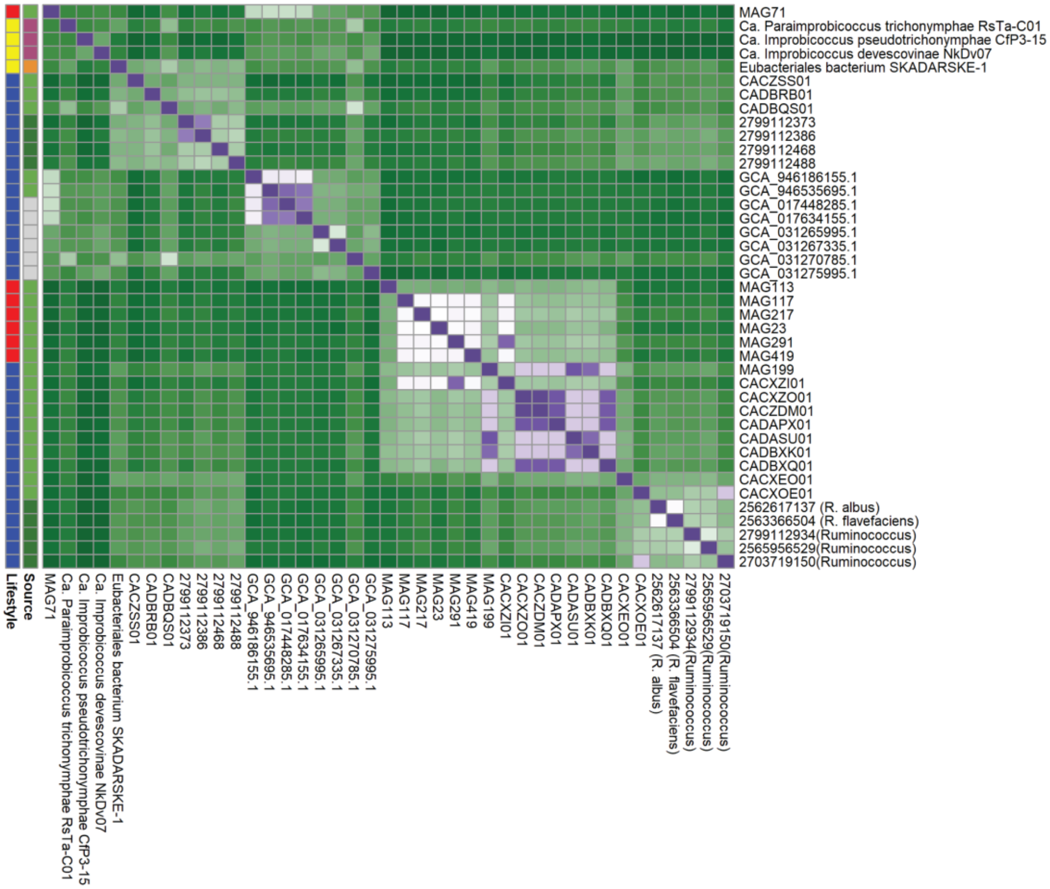


b

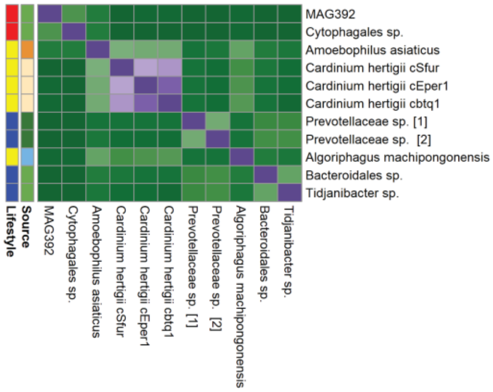


**Figure S4. Amino acid identity.** Amino acid identity (AAI) of the (**a**) Bacteroidota and (**b**) Bacillota candidate symbiont genomes. **a.** Pairwise AAI values between the rumen Bacteroidota symbiont *Candidatus Cillophilus ruminis* (MAG-392) and selected genomes from the Amoebophilaceae family, including *Candidatus Amoebophilus asiaticus* and the previously assembled rumen genome CAMJVU01. MAG-392 shares highest identity with *Ca. A. asiaticus* (AAI = 49.96%), supporting its classification as a novel genus within Amoebophilaceae. **B.** AAI analysis of the Bacillota candidate symbiont MAG-71 (Acutalibacteraceae) relative to known rumen MAGs and

Isolates and members of the small-genome clade identified within the family, including symbionts of termite gut flagellates (*Ca. Improbicoccus pseudotrichonymphae*, *Ca. I. devescovinae*, *Ca. Paraimprobicoccus trichonymphae*). MAG-71 clusters closely with these known and putative symbionts, supporting its placement within a lineage of gut-associated, genome-reduced intracellular bacteria.
